# Involvement of Hepatitis B Core–Related Antigen in Viral Genome Integration in Patients With Prior Hepatitis B Virus Infection

**DOI:** 10.1101/2024.11.21.624775

**Authors:** Tomoya Saito, Rigel Suzuki, Akhinur Rahman, Kento Mori, Samiul Alam Rajib, Nobuhiro Kobayashi, Takaya Ichikawa, Tatsuya Orimo, Tatsuhiko Kakisaka, Lihan Liang, Naganori Nao, Saori Suzuki, Tomokazu Tamura, Yorifumi Satou, Akinobu Taketomi, Takasuke Fukuhara

## Abstract

**Background:** The incidence of non-B non-C hepatocellular carcinoma (NBNC-HCC), which is negative for hepatitis B surface antigen and hepatitis C virus antibodies, is on the rise. Relatively high numbers of NBNC-HCC patients are hepatitis B core antibody (HBcAb) positive, suggesting that previous HBV infection may play a role in NBNC-HCC development, though the exact mechanisms are unclear. This study aimed to investigate whether HBV genomes are integrated into the host genome of HBcAb-positive NBNC-HCC cases and how these integrations may contribute to cancer development and progression.

**Methods:** HBV detection PCR using HBV-specific primers on DNA extracted from HBcAb-positive NBNC-HCC tissue samples was performed. Positive samples were further examined for HBV integration sites using viral DNA–capture sequencing. Additionally, hepatitis B core-related antigen (HBcrAg) serum levels were measured to assess whether they could be predictive for HBV detection PCR results.

**Results:** Among 90 HBcAb-positive NBNC-HCC samples, HBV genome amplification was detected in 18 samples, and elevated HBcrAg levels were associated with the HBV detection PCR results. Seventeen of these samples exhibited HBV integration. The HBV genome was integrated near the *TERT* gene in 7 samples, resulting in significantly increased *TERT* mRNA levels; in the *KMT2B* gene (2 samples); and downstream of *LOC441666* (2 samples).

**Conclusion:** The integration sites we identified in our samples have been previously reported in HBV-related HCC, suggesting that HBV integration may also contribute to hepatocarcinogenesis in HBcAb-positive NBNC-HCC. Furthermore, HBcrAg could serve as a potential, noninvasive marker for detecting HBV integration in these cases.

## Introduction

Liver cancer affects approximately 860,000 people worldwide, ranking sixth in incidence among all cancers. Despite advances in treatment, it continues to have a poor prognosis with around 760,000 deaths annually and is the third leading cause of cancer-related deaths globally. The disease is notably prevalent in developing countries, particularly in Asia, and its incidence is expected to rise in many regions in the future [1].

Hepatocellular carcinoma (HCC) accounts for approximately 90% of all liver cancers. Historically, the majority of HCC cases were attributed to infections by the hepatitis B virus (HBV) or hepatitis C virus (HCV). However, due to the implementation of infection control measures, such as HBV vaccination, the incidence of HBV-related HCC (B-HCC) has declined significantly [2]. Similarly, the advent of antiviral therapies has led to a decrease in HCV-related HCC (C-HCC) [3, 4]. In contrast, the proportion of non-B non-C hepatocellular carcinomas (NBNC-HCC), defined as negative for both hepatitis B surface antigen (HBsAg) and anti-HCV antibodies, has been increasing [5, 6]. The etiology of NBNC-HCC includes factors such as diabetes mellitus, alcohol-related liver disease, and nonalcoholic fatty liver disease (NAFLD), but approximately 40% of cases remain idiopathic [7]. Additionally, patients with NBNC-HCC may have previously had an HBV infection, as indicated by the presence of HBV core antigen antibodies (HBcAb). Notably, the incidence of liver-related diseases is significantly higher in HBcAb-positive vs HBcAb-negative cases, with 73.9% of patients with NAFLD-related or idiopathic HCC being HBcAb-positive [8]. Moreover, HBV reactivation has been observed in some HBcAb-positive individuals upon treatment with immunosuppressive or anticancer drugs, even in the absence of HBsAg or hepatitis B surface antibody (HBsAb) [9]. These findings suggest that NBNC-HCC patients with prior HBV infection (HBcAb positive) may harbor HBV genomes, but it remains unclear whether these viral genomes contribute to carcinogenesis or cancer progression. Furthermore, no effective blood biomarkers have been identified for detecting such cases.

HBV is an enveloped DNA virus with a small 3.2 kbp partially double-stranded genome. HBV enters hepatocytes by binding to the sodium taurocholate co-transporting polypeptide (NTCP). Inside the cell, HBV is converted from relaxed circular DNA (rcDNA) to covalently closed circular DNA (cccDNA) within the nucleus. Viral mRNAs, including the pregenomic RNA (pgRNA), are transcribed from cccDNA. The pgRNA is reverse transcribed into rcDNA within the core particle, forming the nucleocapsid, which associates with HBs antigen and is ultimately released as an infectious particle. A portion of rcDNA is also recycled back into the nucleus [10]. Additionally, double-stranded linear DNA (dslDNA), generated during reverse transcription from pgRNA, can integrate into the host genome via non-homologous end joining [11]. Thus, individuals with a history of HBV infection may have dslDNA integrated into their genome, which could contribute to hepatocarcinogenesis, even when viral replication is suppressed [12]. Furthermore, HBV single-stranded DNA (ssDNA) and spliced variants may also play a role, with pathways such as microhomology-mediated end joining and single-stranded annealing potentially involved, although these mechanisms are not yet fully characterized [13].

HB core–related antigen (HBcrAg) has been reported as a marker for understanding the pathogenesis of chronic hepatitis B [14]. HBcrAg reflects the amount of cccDNA n the liver and is a collective term for hepatitis B envelope antigen (HBeAg), hepatitis B core antigen (HBcAg), and the 22-kDa HBV precore antigen (p22cr). It has been reported that HBcrAg may remain detectable in patients even with undetectable serum HBV DNA or loss of HBsAg [15]. High-sensitivity HBcrAg testing is also considered a therapeutic target for hepatitis B and has been noted as useful in predicting liver carcinogenesis [16]. Additionally, HBcrAg-based gray zone (GZ)–HCC scores have been shown to predict HCC risk better than HBV DNA–based scores in HBeAg-negative patients, making it a valuable tool for clinical management [17]. Based on these reports, we hypothesize that HBcrAg may serve as a marker for detecting high-risk liver carcinogenesis in HBcAb-positive NBNC -HCC and for identifying patients with HBV genome integration.

In this study, we aimed to investigate the presence of HBV genomes in HBcAb-positive NBNC-HCC cases and their potential role in hepatocarcinogenesis and cancer promotion. We analyzed blood and liver tissue samples from 90 patients with HBcAb-positive NBNC-HCC. Using HBV-targeted primers, we identified 18 cases (20%) with amplification of HBV-specific DNA. Most of these cases were positive for HBcrAg, demonstrating superior sensitivity compared to serum HBV DNA testing. These findings suggest that HBcrAg could be a valuable marker for preoperative detection of high-risk cancer without the need for invasive procedures like liver biopsy.

Furthermore, we used next-generation Viral DNA–Capture sequencing Approach (VDCA) to confirm the presence of the HBV genome and identify the insertion sites within the host genome. Interestingly, HBV sequence was predominantly integrated near the oncogene telomerase reverse transcriptase (*TERT*). Elevated levels of *TERT* mRNA were observed in samples where HBV integration was near the *TERT* gene, suggesting a possible role in carcinogenesis. These results indicate that HBV genomes may be integrated into the host genome in some HBcAb-positive NBNC-HCC cases, particularly upstream of the *TERT* gene, and thus contribute to cancer development. Thus, high-sensitivity HBcrAg testing could be a key, non-invasive approach for readily identifying these high-risk patients.

## Materials and Methods

### Patients

We reviewed data from 500 patients with a diagnosis of hepatocellular carcinoma who underwent hepatectomy at the Department of Gastroenterological Surgery I of Hokkaido University Hospital between January 2010 and July 2021. We excluded 367 patients who were hepatitis B surface antigen (HBsAg)–positive, anti-hepatitis C virus (HCV) antibody (HCV-Ab)–positive, and/or hepatitis B core antigen antibody (HBcAb)–negative. An additional 43 patients were excluded who did not have blood or liver specimens available and/or did not have data on HBsAb, HBeAg, and HBeAb levels. Consequently, samples from 90 patients were eligible for this study (Fig.1).

**Figure 1.**
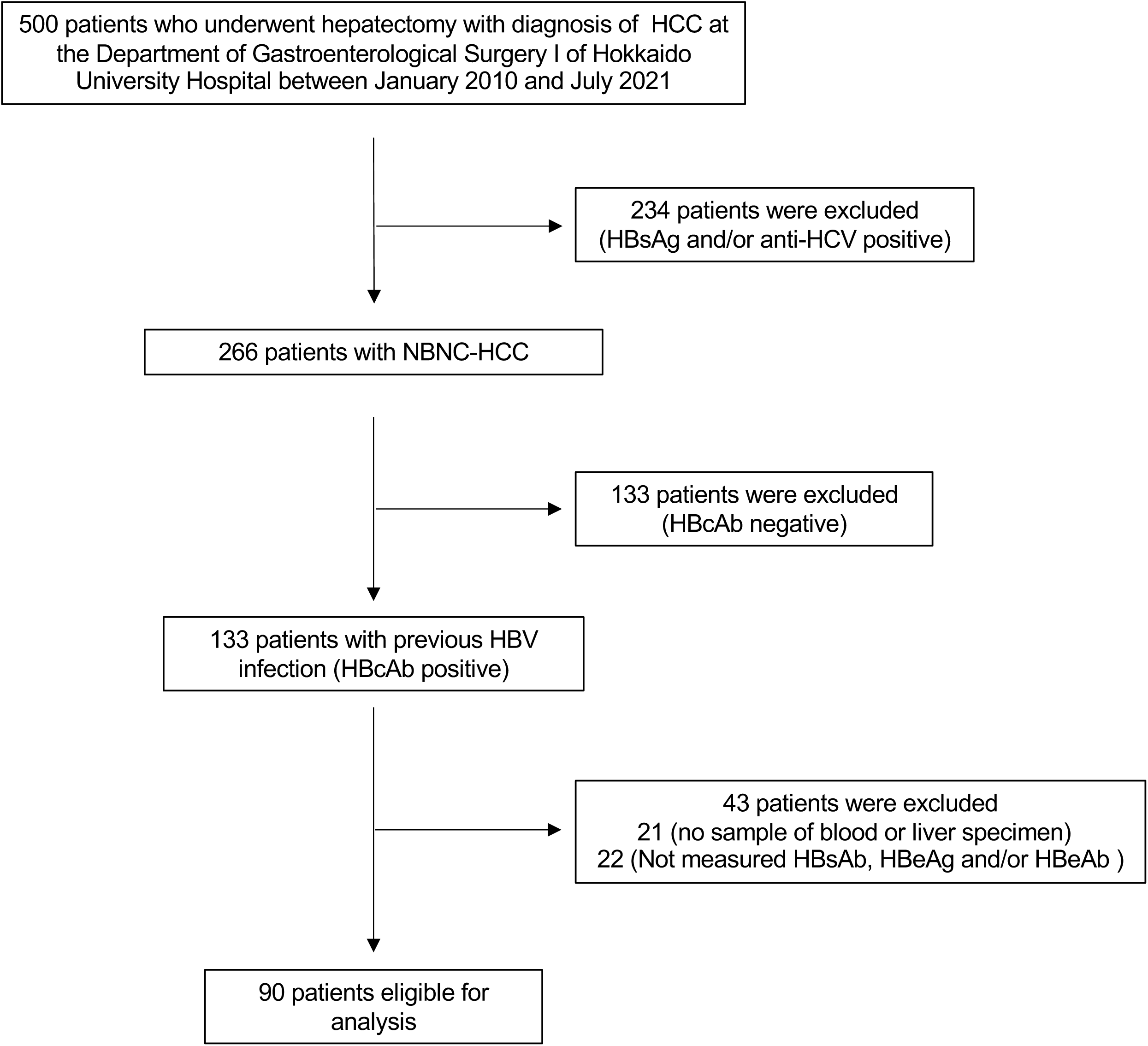
Selection of specimens for analysis. Abbreviations: HCC, hepatocellular carcinoma; HBsAg, Hepatitis B surface antigen; HBV, hepatitis B virus; HCV, hepatitis C virus; HBcAb, hepatitis B core antibody; HBeAg, Hepatitis B e antigen; HBeAb, Hepatitis B e antibody

Liver tissues were obtained at surgery and immediately frozen in liquid nitrogen and stored at -80°C. HBcAb-positive cases were defined as those with prior HBV infection. Frozen tissue specimens and frozen sera stored in the tissue bank were used as specimens. HBV-related HCC (B-HCC) samples were used as positive controls. Ethics approval was obtained from the Hokkaido University Hospital (Ethics application no. 022-0325). This study was conducted in accordance with the principles of the 1996 Declaration of Helsinki. Written informed consent was obtained from all patients.

### Viral DNA Capture-Seq Approach

The DNA extracted from cancerous parts of HBcAb-positive NBNC-HCC was sonicated using Biorupter BR-II (Cat#BR2006A, BMBio; sonication time: 300s; strength of sonication: high) and fragmented to approximately 300-500 bp. Using the NEBNext^®^ UltraTM II DNA library prep kit for Illumina (Cat#E7645S, NEB), the library was further fragmented to 300 bp per the manufacturer’s protocol. A TapeStation (Cat#G2991BA, Agilent Technologies) system was used to confirm that the DNA was fragmented to 300 bp. Next, a probe-based enrichment step was performed as previously reported [18]. Briefly, the multiplexed library was mixed with 53 biotinylated DNA probes (Table S1) targeting HBV genotype C sequences (GenBank accession no. AB246345). Hybridization was performed by incubating the mixtures at 65°C for 4 h. Streptavidin-coated beads were added, and after several washing steps (xGen lockdown reagent, Cat#1072281, IDT), the captured DNA fragments were PCR-amplified with P5 and P7 primers for Illumina sequencing. Enriched DNA libraries were quantified using a TapeStation system and quantitative PCR (GenNext NGS Library Quantification Kit, Code No.NLQ-101, Toyobo). Sequencing was performed using Illumina MiSeq instruments (Cat#SY-410-1003, Illumina) with 2 × 75 bp reads.

### High-throughput sequencing data analysis

Three fastq files—Read 1, Read 2 and Index Read—were obtained from Illumina MiSeq. We first performed a data-cleaning step by using an in-house Perl script (kindly provided by Dr Michi Miura, Imperial College London), which extracts reads with a high Index Read sequencing quality (Phred score >20 in each position of the 8-bp index read). We next removed adaptor sequences from Read 1 and Read 2 followed by a cleaning step to remove reads with too short or with too low of a Phred score as previously described [19]. The cleaned sequencing reads were aligned to a human reference genome (hg19) and HBV (GenBank accession no. AB246345) as a separate chromosome or integrated provirus using the BWA-MEM algorithm [20]. We then used Samtools [20] and Picard (http://broadinstitute.github.io/picard/) for further data processing and cleanup (eg, removal of reads with multiple alignments and duplicated reads).

Details of other experiments can be found in the supplementary material.

## Results

### Detection of HBV genomes in HBcAb-positive NBNC-HCC and utility of serum HBcrAg as a potential marker for HBV genomes in HBcAb-positive NBNC-HCC

A significant number of NBNC-HCC patients have been reported as HBcAb positive [21]. In fact, in our study, 133 of 266 NBNC-HCC samples (50%) were HBcAb-positive, indicating a high prevalence of prior HBV infection among NBNC-HCC patients. To investigate whether HBcAb-positive NBNC-HCC cases contained the HBV genome, we performed HBV DNA detection by PCR. We reviewed the data of 500 patients with a diagnosis of HCC who underwent a hepatectomy at the Hokkaido University Hospital; 367 patients who were HBsAg-positive, HCV antibody-positive, and/or HBcAb-negative were excluded. Patients without available blood or liver specimens or without measurements of HBsAb, HBeAg, and HBeAb levels were also excluded, leaving 90 patients for further analysis (Fig.1).

HBV DNA detection PCR of total DNA extracted from HBcAb-positive NBNC-HCC liver tissue was performed using 5 primer sets specific to HBV genotype C, the most common HBV genotype in Japan [22] (Table S1). Total DNA from B-HCC samples was used as a positive control. HBV genomes were detected in 18 out of 90 samples (Fig.2A).

**Figure 2.**
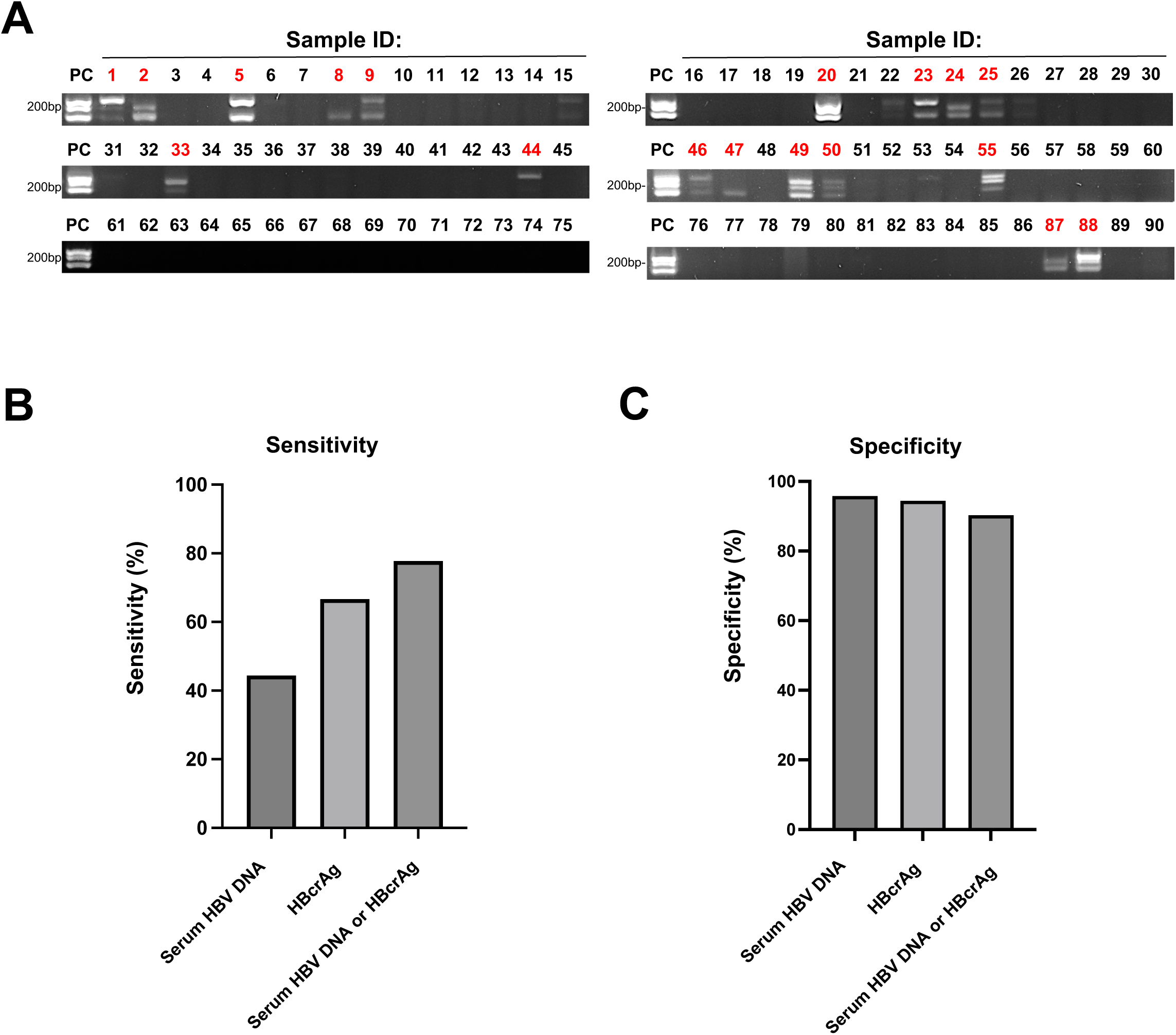
The HBV genome is detected in cancerous areas of HBcAb-positive NBNC-HCC by PCR and HBcrAg is a suitable marker for their identification. **A** DNA was extracted from cancerous parts of HBcAb-positive NBNC-HCC samples, and PCR was performed with 5 HBV-specific primer sets. Eighteen of 90 samples were identified as positive. Positive case numbers are shown in red. DNA extracted from cancerous parts of B-HCC was used as a positive control. **B-C** HBcrAg and HBV DNA were measured in frozen sera from HBcAb-positive NBNC-HCC patients. Then, sensitivity (B) and specificity (C) were determined between the findings from HBV DNA detection PCR vs those from assessing HBcrAg, serum HBV DNA, and HBcrAg or serum HBV DNA.

To assess whether HBV genomes in HBcAb-positive NBNC-HCC could also be detected in blood samples, we examined the association between levels of serum HBV DNA or HBcrAg and the results of our HBV detection PCR in liver samples. Serum HBV DNA is commonly used to detect occult HBV infection [23], while HBcrAg levels correlate with the amount of cccDNA in the liver [24]. Therefore, these serum markers were expected to reflect the results of our HBV detection PCR in the liver samples.

We measured serum HBV DNA levels by real-time PCR and HBcrAg levels by chemiluminescence enzyme immunoassay and assessed the sensitivity and specificity of these methods. The sensitivity was 44.4% for serum HBV DNA, 66.7% for HBcrAg, and 77.8% for either HBcrAg or serum HBV DNA positivity (Fig.2B). The specificity was 95.8% for serum HBV DNA, 94.4% for HBcrAg, and 90.3% for either HBcrAg or serum HBV-DNA, with no significant difference between them (Fig.2C). These results suggest that HBcrAg may be a more effective marker than serum HBV DNA for identifying HBcAb-positive NBNC-HCC cases that contain HBV genomes.

Additionally, 16 of the 90 (17.78%) HBcAb-positive NBNC-HCC cases were positive for HBcrAg, indicating a positive correlation between HBV detection PCR results and HBcrAg levels (p <0.0001) (Table 1).

**Table 1.** Correlation between HBV detection PCR results and HBcrAg.

| Variables |  | HBV detection PCR |  | P value |
| --- | --- | --- | --- | --- |
|  |  | Negative (n=72) | Positive (n=18) |  |
| HBcrAg | Negative(n=74) | 68 | 6 | p<0.0001 |
|  | Positive(n=16) | 4 | 12 |  |

\* HBcrAg levels below a lower quantification limit of 2.1 log<sub>10</sub> U/mL were considered negativity in this study.

### Establishment of the HBV DNA–Capture Sequencing Approach

Our next goal was to establish a comprehensive method to identify the integration sites of the HBV genome in the host genome of the 18 HBcAb-positive NBNC-HCC samples we identified as containing HBV (Fig.2A). VDCA is a sensitive and comprehensive method for analyzing the insertion sites of viral genomes in the host genome [18]. Conventional next-generation sequencing (NGS) analysis has low sensitivity for detecting HBV genome insertion sites, requiring a large number of reads that results in high costs. In VDCA, DNA extracted from tissue is made into a library and then annealed with biotin-labeled probes specific to the viral (in this case, HBV) genome. The probes are then captured using streptavidin beads, and the host genome sequence which the viral genome is inserted among is then enriched for. The reads obtained using this method include chimeric reads of host and viral genomes, allowing the integration sites to be determined by NGS analysis (Fig.3A). This method has been used to identify the integration sites of human T-cell leukemia virus type 1 and bovine leukemia virus [25, 26].

**Figure 3.**
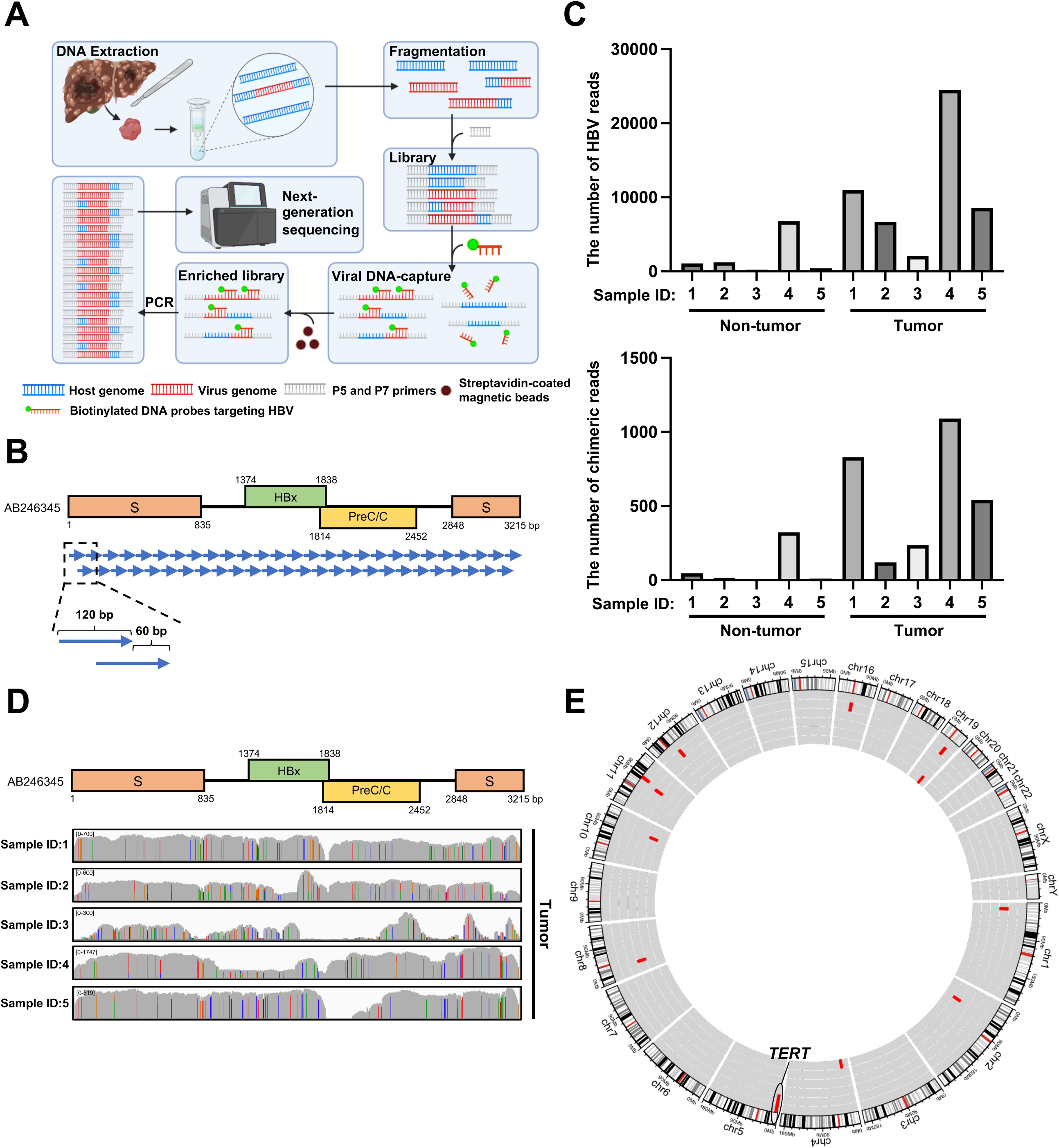
The identification of HBV integration sites using Viral DNA–Capture Sequencing Approach. **A** Experimental workflow of HBV DNA–Capture Sequence Approach. A DNA-probe-based enrichment step was introduced to increase the detection efficiency of HBV-host chimeric reads. **B** Design of the 53 HBV-specific probes, 120 bp in length and with 60 bp tiling. S, PreS1/PreS2/S region; PreC/C, Pre-core/core region; HBx, HBx region. **C** The number of HBV-reads (upper panel) and chimeric-reads (lower panel) were detected by Viral DNA-Capture Sequencing Approach using tumor and non-tumor of 5 B-HCCs. **D** The upper panel shows a schematic of the HBV genome structure. The lower panel shows where reads from the tumor tissue of 5 patients with B-HCC mapped on the HBV genome. **E** Visualization of HBV integration breakpoints in the host DNA of B-HCC samples. Each red bar represents the HBV integration breakpoints at a particular locus in chromosomes of the human genome (hg19). Each inner circle “track” represents the different samples. Chr, chromosome.

In this study, we designed 53 biotin-labeled probes (each 120 bp long, tiling every 60 bp) targeting the HBV genome (GenBank accession number: AB246345) (Fig.3B). To verify the accuracy of these probes, we performed VDCA on total DNA extracted from both cancerous and non-cancerous liver tissues of five patients with B-HCC. We observed that the number of HBV and chimeric reads was higher in cancerous than non-cancerous tissues (Fig.3C). This is because in cancerous tissue, cells with the HBV genome integrated into the host genome increase clonally. The HBV reads we identified in the cancerous tissues indeed mapped to the reference HBV genome and covered almost the entire viral genome (Fig.3D). These results indicated that the designed probes could comprehensively capture HBV genomic sequence. Next, we identified the HBV integration sites in the cancerous tissues. Consistent with previous reports [27], we found that in 2 of the 5 cases the HBV genome was inserted into the *TERT* gene (Fig.3E). These results demonstrated the suitability of the VDCA method for identifying HBV integration sites.

### Identification of HBV Integration Sites in HBcAb-Positive NBNC-HCC

Next, we used VDCA to examine whether HBV integration sites were present in the 18 HBcAb-positive NBNC-HCC samples that we identified as containing HBV genome. Based on the results in Fig.3C, which showed more chimeric reads in cancerous tissues compared to non-cancerous tissues, we focused our analysis on the cancerous tissues collected from these patients. Among the 18 samples, one had <500 HBV reads and <50 chimeric reads, and no integration sites were identified (Fig.4A). Among the remaining 17 samples, the HBV genome was integrated into the *TERT* gene in 7 cases, the *KMT2B* gene in 2 cases, and the downstream of *LOC441666* region in 2 cases (Fig.4B). The other integration sites identified are listed in Table 2. Of note, many of the samples with HBV genome integration in the *TERT* gene had no other identified integration sites. In contrast, samples without HBV integration in *TERT* exhibited multiple integration sites (Table 2). These results suggest that hepatocarcinogenesis in NBNC-HCC may be driven by the insertion of HBV genes into cancer-related driver genes such as *TERT* or into various other regions of the human genome.

**Figure 4.**
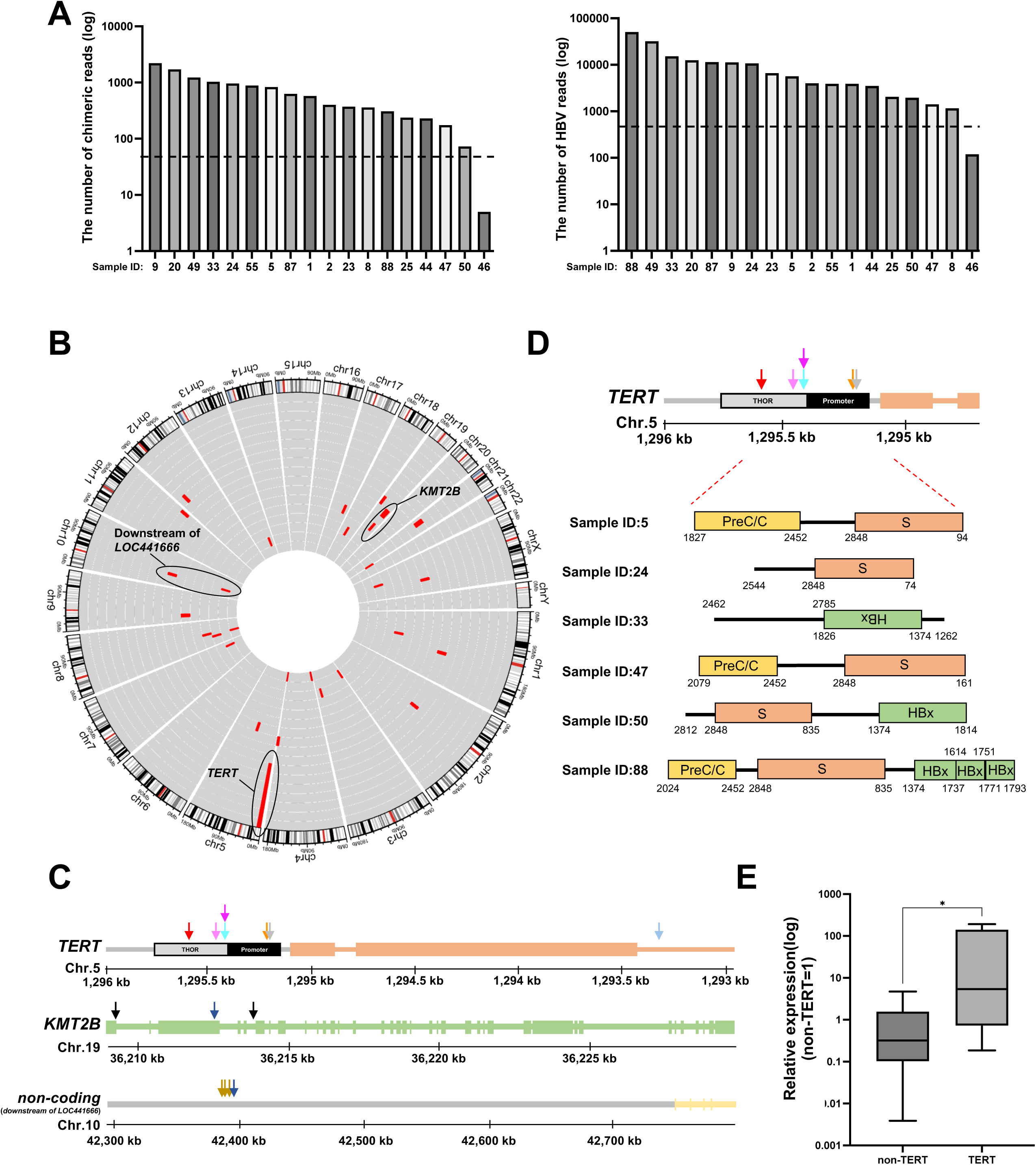
The identification of HBV integration breakpoints in HBcAb-positive NBNC-HCC. **A** The number of HBV reads (left panel) and chimeric reads (right panel) detected by Viral DNA–Capture Sequencing Approach in 18 HBcAb-positive NBNC-HCC liver tissue samples. The dotted line in the left panel is at a value of 50 and the dotted line in the right panel is at a value of 500. **B** Visualization of HBV integration breakpoints in the host DNA of HBcAb-positive NBNC-HCC samples. Each red bar represents the HBV integration breakpoints at a particular locus in chromosomes of the human genome (hg19). Each inner circle “track” represents the different samples. **C** Mapping of HBV breakpoint integration sites. The HBV breakpoint sites on the recurrently affected genes *TERT, KMT2B* and region downstream of *LOC441666* mapped to the human hg19 reference sequence. Each arrow represents the location of an HBV breakpoint identified in a clinical sample. Arrows of the same color indicate HBV breakpoints found in the same specimen. Boxes represent exons. Chr, chromosome. **D** Regions of the HBV genome for each of the 6 cases in which the HBV genome was integrated into the promoter or THOR of *TERT*. S, PreS1/PreS2/S region; PreC/C, Pre-core/core region; HBx, HBx region. **E** Ratio of expression of *TERT* mRNA levels in HBcAb-positive NBNC-HCC samples with HBV genome integration into or near the *TERT* gene (n=6) vs those with integration in other regions (n=10. RNA was extracted from these samples, and the expression level of the *TERT* gene was quantified relative to *GAPDH* using theΔΔCt method. Significant difference was found. *P < 0.05 as determined by Mann-Whitney U test.

**Table 2.** HBV integration sites in the HBcAb-positive NBNC-HCC samples positive for HBV by detection PCR.

| Sample | chr | Annotation | Symbol | Gene Name |
| --- | --- | --- | --- | --- |
| 1 | chr1 | Intron | ANKRD20A12P | ankyrin repeat domain 20 family member A12, pseudogene |
| 1 | chr1 | Intron | NA | NA |
| 1 | chr11 | Intron | NTM | neurotrimin |
| 1 | chr2 | Intron | LOC442028 | uncharacterized LOC442028 |
| 1 | chr21 | Intergenic | MIR3648-1 | microRNA 3648-1 |
| 1 | chr21 | Intergenic | ANKRD20A11P | ankyrin repeat domain 20 family member A11, pseudogene |
| 2 | chr5 | Intron | XRCC4 | X-ray repair cross complementing 4 |
| 5 | chr5 | Promoter | TERT | telomerase reverse transcriptase |
| 8 | chr18 | Intron | CCDC178 | coiled-coil domain containing 178 |
| 8 | chr3 | Intergenic | ZBBX | zinc finger B-box domain containing |
| 8 | chr8 | Intergenic | LINC01299 | long intergenic non-protein coding RNA 1299 |
| 9 | chr21 | Intergenic | PRMT2 | protein arginine methyltransferase 2 |
| 9 | chr3 | Intergenic | ZBBX | zinc finger B-box domain containing |
| 9 | chr8 | Intergenic | LINC00293 | long intergenic non-protein coding RNA 293 |
| 9 | chrX | Intergenic | ZC4H2 | zinc finger C4H2-type containing |
| 20 | chr10 | Intergenic | LOC441666 | zinc finger protein 91 pseudogene |
| 20 | chr10 | Intergenic | LOC441666 | zinc finger protein 91 pseudogene |
| 20 | chr10 | Intergenic | LOC441666 | zinc finger protein 91 pseudogene |
| 20 | chr13 | Intergenic | MIR4500 | microRNA 4500 |
| 20 | chr3 | Intergenic | CNTN6 | contactin 6 |
| 20 | chr7 | Intergenic | GRM8 | glutamate metabotropic receptor 8 |
| 23 | chr4 | Intergenic | ZNF595 | zinc finger protein 595 |
| 23 | chr4 | Intergenic | DUX4L8 | double homeobox 4 like 8 (pseudogene) |
| 23 | chr8 | Intergenic | XKR4 | XK related 4 |
| 24 | chr12 | Intergenic | AEBP2 | AE binding protein 2 |
| 24 | chr5 | Intron | TERT | telomerase reverse transcriptase |
| 25 | chr5 | Promoter | TERT | telomerase reverse transcriptase |
| 33 | chr5 | Promoter | TERT | telomerase reverse transcriptase |
| 44 | chr19 | Intron | SBNO2 | strawberry notch homolog 2 |
| 47 | chr5 | Promoter | TERT | telomerase reverse transcriptase |
| 49 | chr10 | Intergenic | LOC441666 | zinc finger protein 91 pseudogene |
| 49 | chr19 | Exon | KMT2B | lysine methyltransferase 2B |
| 49 | chr19 | Intron | CBLC | Cbl proto-oncogene C |
| 49 | chr4 | Intron | SORBS2 | sorbin and SH3 domain containing 2 |
| 49 | chrX | Intergenic | RPA4 | replication protein A4 |
| 50 | chr5 | Promoter | TERT | telomerase reverse transcriptase |
| 55 | chr19 | Intron | KMT2B | lysine methyltransferase 2B |
| 55 | chr17 | Intron | BCAS3 | BCAS3 microtubule associated cell migration factor |
| 55 | chr19 | Intron | KMT2B | lysine methyltransferase 2B |
| 55 | chr9 | Promoter | IFNA14 | interferon alpha 14 |
| 55 | chr9 | Intergenic | LINC03106 | long intergenic non-protein coding RNA 3106 |
| 87 | chr5 | Promoter | TERT | telomerase reverse transcriptase |
| 88 | chr1 | Intron | PRMT6 | protein arginine methyltransferase 6 |

Further analysis of the HBV integration sites in the host genome revealed that HBV was inserted into the promoter region or TERT hypermethylated tumor region (THOR) of the *TERT* gene. THOR is a 433 bp genomic region containing 52 CpG sites located immediately upstream of the *TERT* core promoter. The methylation status of this region has been reported to influence the expression level of *TERT* [28]. Moreover, 2 samples had HBV integrated into intron 1, exon 3, and intron 5 of the *KMT2B* gene. *KMT2B* encodes a histone methyltransferase that regulates chromatin structure and gene transcription involved in carcinogenesis [29]. Additionally, we identified HBV integration in a non-coding region downstream of *LOC441666* (Fig.4C). According to Repeat Masker annotation, the HBV integration sites downstream of *LOC441666* were in an RNA repeat sequence called Human Satellite II (HSATII), which has been implicated in the malignant transformation of pancreatic cancer [30]. These results suggest HBV integration forms human-virus chimeric genes that may affect the expression of downstream or neighboring genes, potentially leading to hepatocellular carcinoma.

Next, we investigated which regions of HBV were integrated into the host genome. In this study, we performed NGS analysis of short reads. As a result, we were able to identify sequences at both ends of the HBV fragments that had integrated into the host genome but could not accurately identify the sequences in between. The integration near the TERT gene was confirmed in six cases, indicating the need for further analysis. To address this limitation, we designed primers targeting the HBV integration regions in the promoter or THOR of *TERT* and performed sequencing PCR of the 6 samples with HBV integrated in these regions. We found that 5 of the 6 samples contained 2848-3200bp and 1-74 bp of the HBV PreS1/PreS2/S region. Additionally, 3 samples contained 1374-1751 bp and 1771-1793 bp of the HBx region and/or 2079-2452 bp of the PreC/C region (Fig. 4D).

We hypothesized that the HBV DNA integration we observed in the promoter or THOR of *TERT* might affect *TERT* expression levels. To test this, we compared *TERT* expression using RT-qPCR in the 7 cases of HBcAb-positive NBNC-HCC we identified with HBV genome integration in the promoter or THOR of *TERT* vs the 10 cases without. We found that the cases with HBV genome integration near the *TERT* gene exhibited significantly higher *TERT* mRNA expression compared to those without (p = 0.0136) (Fig.4E), suggesting that integration of the HBV genome in this region increases *TERT* mRNA expression.

## Discussion

In this study, we detected HBV genomes in 18 out of 90 samples of HBcAb-positive NBNC-HCC (Fig.2A). After applying VDCA to these samples, we found that HBV genomes were integrated into the host genome in 17 of the 18 samples (Fig.4C), suggesting that approximately 18.9% (17 out of 90 samples) of HBcAb-positive NBNC-HCC patients exhibit HBV integration. The integration hotspots included the promoter and THOR of *TERT*, the *KMT2B* gene, and the downstream regions of the *LOC441666* gene (Fig.4C). The HBV genome was most frequently integrated in the promoter or THOR of *TERT* and was associated with increased expression of *TERT* (Fig.4C and 4E). Both *TERT* and *KMT2B* have been previously identified as HBV integration sites in B-HCC [31-33]. This suggests that some HBcAb-positive NBNC-HCC patients may share characteristics with B-HCC, despite their classification as NBNC-HCC. Additionally, we found that serum HBV DNA as well as HBcrAg could be used to detect HBV genome integration in HBcAb-positive NBNC-HCC cases (Fig.2B and 2C). Various methods, such as *Alu*-PCR and whole genome sequencing (WGS), have been used to analyze HBV integration in B-HCC samples [33, 34]. The *Alu*-PCR method uses HBV-specific primers and primers targeting *Alu* sequences, which are abundant in the human genome, followed by Sanger sequencing to identify integration sites.

However, *Alu*-PCR is limited in its ability to detect integration sites far from *Alu* sequences, making comprehensive analysis difficult [35]. WGS can provide a complete genome analysis but is costly and impractical for large-scale studies [27]. In contrast, VDCA enriches the regions of the host genome that contain HBV sequence using HBV-specific probes, followed by NGS, making it a cost-effective alternative to WGS that still allows for comprehensive analysis. Additionally, VDCA can be performed using either short-or long-read sequencing. While short-read sequencing may not capture the full sequence of integrated viral genomes, this limitation can be addressed, as we did, by reanalyzing the regions around the integration site using PCR (Fig.4D). For more detailed analyses or smaller sample sizes, long-read sequencing may be more appropriate.

Several studies have identified HBV integration sites in B-HCC patients. Sung et al. reported HBV integration at sites such as *TERT, KMT2B, CCNE1, SENP5, ROCK1*, and *FN1*, with about 40% of HBV genome breakpoints occurring within a 1,800 bp region containing the viral enhancer, X gene, and core gene [33]. Similarly, Jiang et al. found HBV integration into *KMT2B, FN1, CTDSPL2*, and *LRP1B*, with the viral integration frequently encompassing the HBV direct repeat sequences DR1 and DR2, which are critical for HBV DNA replication and genome circularization [36]. In contrast, there are few reports investigating potential HBV integration sites in NBNC-HCC. Tamori et al. identified HBV integration in 2 out of 14 NBNC-HCC cases using Southern blot analysis [37]. Another study found HBV integration in 3 of 21 HBsAg-negative HCC cases, with one case showing integration in *TERT* [38]. Tatsuno et al. reported HBV integration in 11 of 73 HCV-negative and HBV-previously infected HCC cases (15.0%), with HBV integration in *TERT* (5.4%) and *KMT2B* (2.7%) [39]. These results are consistent with our findings of integration in or near *TERT* and *KMT2B* and downstream of *LOC441666* (Fig.4C). In our study, the sequences of HBV genome integrated into the promoter or THOR of *TERT* were mainly from the PreS1/PreS2/S region, but also in the X and PreC/C region (Fig.4D), which are commonly associated with B-HCC [40]. Mutations and hypermethylation in the *TERT* promoter and the THOR have been linked to increased *TERT* expression [41]. *TERT* is typically transcriptionally active only in early embryogenesis and highly proliferative cells, but it is reactivated in most cancers, contributing to carcinogenesis by elongating telomeres [42]. *KMT2B* may also play a role in oncogenesis when the HBV genome integrates into exon 3 or intron 5 of *KMT2B*, leading to increased mRNA expression [43]. HBV integration near *LOC441666* has also been reported [44], though its role in hepatocarcinogenesis remains unclear. According to the RepeatMasker annotation, the region downstream of *LOC441666* where we identified HBV integration is located near a repeating sequence known as Human Satellite II (HSATII), which is abnormally expressed in pancreatic cancer and associated with epithelial-mesenchymal transition, enhancing malignancy [30]. Therefore, HBV insertion near these genes may also affect their expression, potentially contributing to hepatocarcinogenesis in HBcAb-positive NBNC-HCC.

Among the 7 cases with *TERT* integration, 6 cases showed no other HBV integrations, whereas cases without *TERT* integration exhibited multiple HBV integration sites (Table 2). This suggests that HBV integration–induced hepatocarcinogenesis may follow 2 distinct mechanisms: one involving strong cancer-related genes like *TERT*, where mutations or altered expression significantly increase cancer risk; the other involving the accumulation of multiple mutations across the host genome, eventually leading to cancer. Previous work has found an association between the number of HBV integrations and early-onset hepatocarcinogenesis [38].

We investigated whether a biomarker that didn’t require biopsy or surgery could be used to help identify patients with HBV genome integration. HBcrAg reflects the amount of cccDNA in the liver and is associated with HCC development [45]. High serum HBcrAg levels (≥10,000 U/mL) have been used to stratify HCC risk in HBeAg-negative patients and to optimize clinical management [46]. However, no studies have linked HBcrAg with HBV integration in HBcAb-positive NBNC-HCC. In this study, we found a positive correlation between HBcrAg positivity and HBV DNA detection in cancer tissues (Table 1), suggesting that HBcrAg could serve as a readily test-able, noninvasive marker for HBV integration in HBcAb-positive NBNC-HCC. Furthermore, HBcrAg showed better sensitivity and specificity compared to serum HBV DNA.

In conclusion, HBV integration into the host genome may contribute to carcinogenesis in some HBcAb-positive NBNC-HCC cases. Measuring HBcrAg levels may facilitate early detection and treatment. However, this study has several limitations. First, it was a single-institution, retrospective study, which may introduce selection bias. Additionally, it remains unclear whether HBcrAg positivity is a cause or consequence of cancer development. Future prospective, multicenter studies are needed. Finally, since NBNC-HCC is defined as HCC that is negative for both HBsAg and HCV antibodies, HBcrAg negativity might be better to include in this definition, as HBcrAg-positive and HBcAb-positive NBNC-HCC are similar to B-HCC.

## Author contributions

Conceptualization: AT and TF; Formal analysis: AR and KM; Funding acquisition: RS, AT and TF; Investigation: TS, RS, AR, KM, SR, NK, TI, LL, NN, SS and TT; Methodology: AR, SR and YS; Project administration: RS and TF; Resources: TS, NK, TO, TK and AT; Software: AR, KM, SR and YS; Supervision: TF; Visualization: TS, RS, KM, LL and TF; Writing: TS, RS, and TF.

## Competing financial interests

The authors declare no competing financial interests.

## Acknowledgements

We thank H. Kubo, M. Tetsuka, and S. Shimamura for their secretarial work and M. Hommura, N. Tachibana, K. Oyama, H. Maruyama, A. Imajo, H. Azuma, Y. Saito and N. Kobayashi for their technical assistance. We also thank Jenna M. Gaska for English proofreading. This work was supported by the Ministry of Health, Labour and Welfare of Japan and the Japan Agency for Medical Research and Development (JP23fk0410052 to Y.S. JP23fk0310524h0002 to A.T. JP23fk0310523h0002, JP23fk0310524h0002, JP23fk0310501h0002, JP23fk0310506h0002, JP23fk0310513h0002 to T.F.), the Japan Society for the Promotion of Science KAKENHI (JP20K22951 to R.S.).

## Supplementary Methods

### DNA extraction

The cancerous parts of B-HCC and HBcAb-positive NBNC-HCC samples that had been stored at -80°C were cut down to 25 mg. Then, these samples were homogenized with a Power Masher II (Cat#891300, NIP), and DNA was extracted using the DNeasy^®^ Blood & Tissue Kit (Cat. No. 69504, QIAGEN) according to the manufacturer’s protocol.

### PCR

HBV sequences in the DNA extracted from HBcAb-positive NBNC-HCC samples were amplified by PCR using a mixture of 5 HBV-specific primer sets (Table S1) and PrimeSTAR^®^ GXL DNA Polymerase (Cat#R050A, Takara Bio) according to the manufacturer’s protocol. Cancerous parts of B-HCC were used as positive control.

### Laboratory assay

HBcrAg and serum HBV DNA levels were measured at SRL Inc. using serum collected prior to the start of surgery from patients with HBcAb-positive NBNC-HCC. Other laboratory data were obtained at the time of the initial outpatient visit or admission. HBcrAg levels below a lower quantification limit of 2.1 log_10_ U/mL were considered negativity in this study. Serum HBV DNA levels below the detection sensitivity were considered negative in this study.

### Integration site analysis using the viral DNA–capture sequencing approach data

To analyze HBV integration sites, we aligned the cleaned fastq files to the reference genome (hg19) containing all human chromosomes (chr 1–22, X, Y and M) and a genotype C HBV reference genome (GenBank accession no. AB246345). In this study, we used an in-house Python script developed to search for HTLV-1 Provirus [1]. This script extracted virus-host reads and generated a comprehensive list of all virus-host reads in the sample. During DNA library preparation, random ligations between viral and host DNAs were generated, resulting in one-sided virus-host reads at either the 5′ or 3′ ends. Consequently, viral integration sites were defined based on the presence of virus-host reads at the 5′ or 3′ end of the provirus, with a mapping depth threshold of five or more virus-host reads. Then, the locations of the integration sites within the human genome were determined by the host-virus junctions present within the sequencing reads. In cases where two host-virus junctions were found in proximity within the human genome, the integration site was defined as located between the region flanked by these junctions. Alternatively, when the host-virus junctions were either distantly located or only one junction was identified, the integration site was defined as occurring at the boundary of the host-virus junction. All mapped regions were visualized and confirmed using the Gviz package in R [2]. During this process, we identified and removed integration sites suspected to be from cross-sample contamination.

### Data and Code Availability

The bam files analyzed from all patients’ samples have been deposited at NCBI BioProject (accession: PRJNA1185153).

### Verifying the HBV sequence integrated into the host genome

DNA extracted from tissue samples of HBcAb-positive NBNC-HCC in which integration of the HBV genome was observed near the *TERT* gene was amplified by PCR using the primers listed in Table S1 and KOD One^®^ PCR Master Mix Blue (Cat#KMM-201, TOYOBO) according to the manufacturer’s protocol. To eliminate nonspecific PCR products, nested PCR was performed by repeating the PCR twice, followed by electrophoresis on an agarose gel. Bands larger than 1000 bp were considered likely to contain HBV integration. These bands were excised from the agarose gel, and gel extraction was performed using the FastGene™ Gel / PCR Extraction Kit (Cat#FG-91302, Nippon Genetics) according to the manufacturer’s instructions. The extracted products were then subjected to sequence analysis using the BigDye™ Terminator v3.1 Cycle Sequencing Kit (Cat#4337455, Thermo Fisher Scientific) following the manufacturer’s protocol, and sequencing was carried out on a SeqStudio Genetic Analyzer (Thermo Fisher Scientific).

### Reverse transcription-quantitative PCR (RT-qPCR)

From the frozen liver tissue samples described above for patients with HBcAb-positive NBNC-HCC, 25 mg was used for RNA extraction. The tissue was homogenized with a Power Masher II (Cat#891300, NIP) and suspended in 500 µL of lysis buffer containing 1% 2-mercaptoethanol from the PureLink^TM^ RNA Mini Kit (Cat#12183018A, Thermo Fisher Scientific). The suspension was then centrifuged at 1500 g for 5 min at room temperature, and 350 µl of the supernatant was used for RNA extraction per the manufacturer’s protocol. The extracted RNA was then transferred to a QuantStudio 5 Real-Time PCR System (Thermo Fisher Scientific) using the One Step TB Green® PrimeScript™ PLUS RT-PCR Kit (Perfect Real Time) (Cat#RR096A, Takara Bio). mRNA levels were quantified using primers for *TERT* and *GAPDH* (Table S1). *GAPDH* was used as a control and the relative expression of *TERT* was calculated using the ΔΔCt method.

---

**Table S1.**
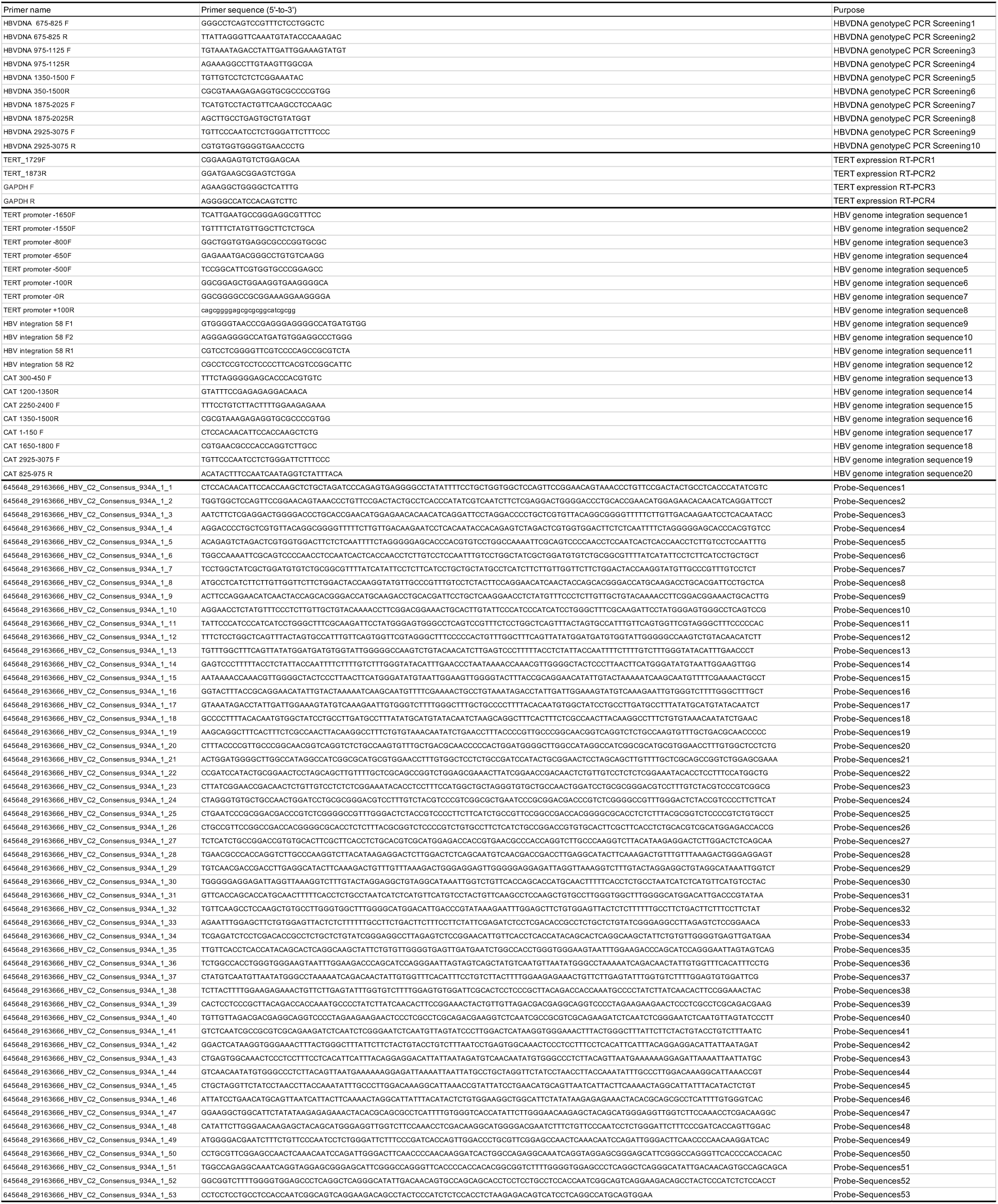
Primers and probes used in this study.

## Notes

### Competing Interest Statement

The authors have declared no competing interest.

